# Ligand promiscuity in the tryptophan repressor – from structural understanding towards rational design

**DOI:** 10.1101/2020.01.20.912972

**Authors:** Andre C. Stiel, Sooruban Shanmugaratnam, Ole Herud-Sikimic, Gerd Jürgens, Birte Höcker

## Abstract

Receptors that promiscuously bind a range of ligands provide insights into how nature mediates affinity and biological functioning. Moreover, such receptors provide vantage points for the rational design of specific binding for biotechnological applications. Here we describe the molecular details of the ligand binding promiscuity of the well-known tryptophan repressor TrpR. We elucidated high-resolution structures of TrpR bound to the co-repressors 5-methyl-tryptophan and 5-methyl-tryptamine as well as the pseudo-repressors indole-3-propionic and indole-3-acetic acid. Furthermore, using isothermal titration calorimetry we procure the corresponding thermodynamic parameters. Together this data provides molecular explanations for the strongly varied affinities and biological effects of the ligands providing insights into how nature shapes specificity and affinity. Beyond this we use these insights to exemplarily showcase knowledge-based design of TrpR by swapping its specificity from its native ligand tryptophan to indole-3-acetic acid. Finally, we elucidate the structures of the variant bound to indole-3-acetic and indole-3-propionic acid to retrace our design rationale.

## Introduction

Promiscuity in ligand binding is a common phenomenon. Molecules related to the native ligand bind to its receptor with decreased or equal affinity and can activate or block native functionality. A molecular understanding of how different ligands bind to the same receptor provides insight into how nature tunes affinity and modulates biological functioning^1^. In addition, such promiscuity is a vantage point for rationally designing protein specificity to molecules of interest^2-4^. For the tryptophan repressor (TrpR) such promiscuity is firmly established: Next to its native ligand tryptophan (TRP) TrpR possesses a number of co-repressors and pseudo-repressors (Figure 1A) that share the indole-group but have different substituents^5^ resulting in different affinities of the ligand to TrpR and of TrpR to its operator DNA^6^. Among those are molecules of high interest for phytochemical applications, like the ubiquitous plant hormone Indole-3-acetic acid (IAA, Auxin) or Indole-3-butyric acid. TrpR is an obligatory symmetric homo-dimer that independently binds two molecules of TRP^7^ (Figure 1B). The ligand binding pockets are situated between the central core and the E helices of the DNA-interacting reading head motifs. TRP is firmly bound by anchoring its amino-group to the backbone of the protein core while the carboxyl-group interacts with the guanidino-group of Arg_84_ in helix E (Figure 1C). This configuration stabilizes the reading-head and poises the arginine for operator-DNA interaction^8,9^ (Suppl. Figure S3A). In contrast, the Arg_84_ side-chain in the apo-form collapses into the binding pocket leaving helix E distorted and preventing DNA interaction^9^. Binding of non-native ligands in this pocket has been investigated in detail with biophysical methods but only few conclusions could be drawn on a molecular level^10,11^. Here we present structural information of the TrpR protein bound to a set of co- and pseudo-repressors (Figure 1) providing insight into the molecular details of ligand binding and its influence on DNA interaction. Moreover, we complement the structures with affinity data from isothermal titration calorimetry (ITC) measurements providing thermodynamic parameters (Table 1, Supp. Figure S1). Exemplarily, we harness this detailed understanding of the TrpR binding pocket to rationally swap its specificity from the native ligand TRP to indole-3-acetic acid by a single mutation. Finally, we retrace the effects using crystal structures of the mutant to explain how specificity can be designed.

**TABLE 1:**
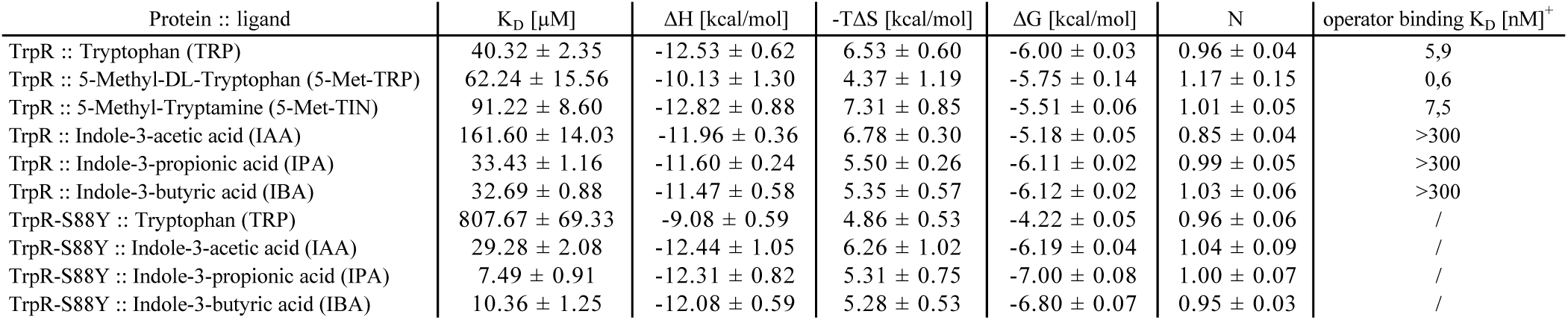
ITC derived thermodynamic properties of TrpR and TrpR-S88Y bound to studied ligands and K_D_s for binding to operator DNA (^+^ from Marmorstein et al., 1989). Data is shown as the mean and standard deviation of n = 3 (TrpR :: TRP and TrpR :: 5-Met-TRP, n = 4) biologically independent replicates (separate purifications of the protein).

| Protein :: ligand | $K_D$ [μM] | $\Delta H$ [kcal/mol] | $-T\Delta S$ [kcal/mol] | $\Delta G$ [kcal/mol] | N | operator binding $K_D$ [nM]<sup>+</sup> |
| --- | --- | --- | --- | --- | --- | --- |
| TrpR :: Tryptophan (TRP) | 40.32 ± 2.35 | -12.53 ± 0.62 | 6.53 ± 0.60 | -6.00 ± 0.03 | 0.96 ± 0.04 | 5,9 |
| TrpR :: 5-Methyl-DL-Tryptophan (5-Met-TRP) | 62.24 ± 15.56 | -10.13 ± 1.30 | 4.37 ± 1.19 | -5.75 ± 0.14 | 1.17 ± 0.15 | 0,6 |
| TrpR :: 5-Methyl-Tryptamine (5-Met-TIN) | 91.22 ± 8.60 | -12.82 ± 0.88 | 7.31 ± 0.85 | -5.51 ± 0.06 | 1.01 ± 0.05 | 7,5 |
| TrpR :: Indole-3-acetic acid (IAA) | 161.60 ± 14.03 | -11.96 ± 0.36 | 6.78 ± 0.30 | -5.18 ± 0.05 | 0.85 ± 0.04 | >300 |
| TrpR :: Indole-3-propionic acid (IPA) | 33.43 ± 1.16 | -11.60 ± 0.24 | 5.50 ± 0.26 | -6.11 ± 0.02 | 0.99 ± 0.05 | >300 |
| TrpR :: Indole-3-butyric acid (IBA) | 32.69 ± 0.88 | -11.47 ± 0.58 | 5.35 ± 0.57 | -6.12 ± 0.02 | 1.03 ± 0.06 | >300 |
| TrpR-S88Y :: Tryptophan (TRP) | 807.67 ± 69.33 | -9.08 ± 0.59 | 4.86 ± 0.53 | -4.22 ± 0.05 | 0.96 ± 0.06 | / |
| TrpR-S88Y :: Indole-3-acetic acid (IAA) | 29.28 ± 2.08 | -12.44 ± 1.05 | 6.26 ± 1.02 | -6.19 ± 0.04 | 1.04 ± 0.09 | / |
| TrpR-S88Y :: Indole-3-propionic acid (IPA) | 7.49 ± 0.91 | -12.31 ± 0.82 | 5.31 ± 0.75 | -7.00 ± 0.08 | 1.00 ± 0.07 | / |
| TrpR-S88Y :: Indole-3-butyric acid (IBA) | 10.36 ± 1.25 | -12.08 ± 0.59 | 5.28 ± 0.53 | -6.80 ± 0.07 | 0.95 ± 0.03 | / |

**FIGURE 1:**
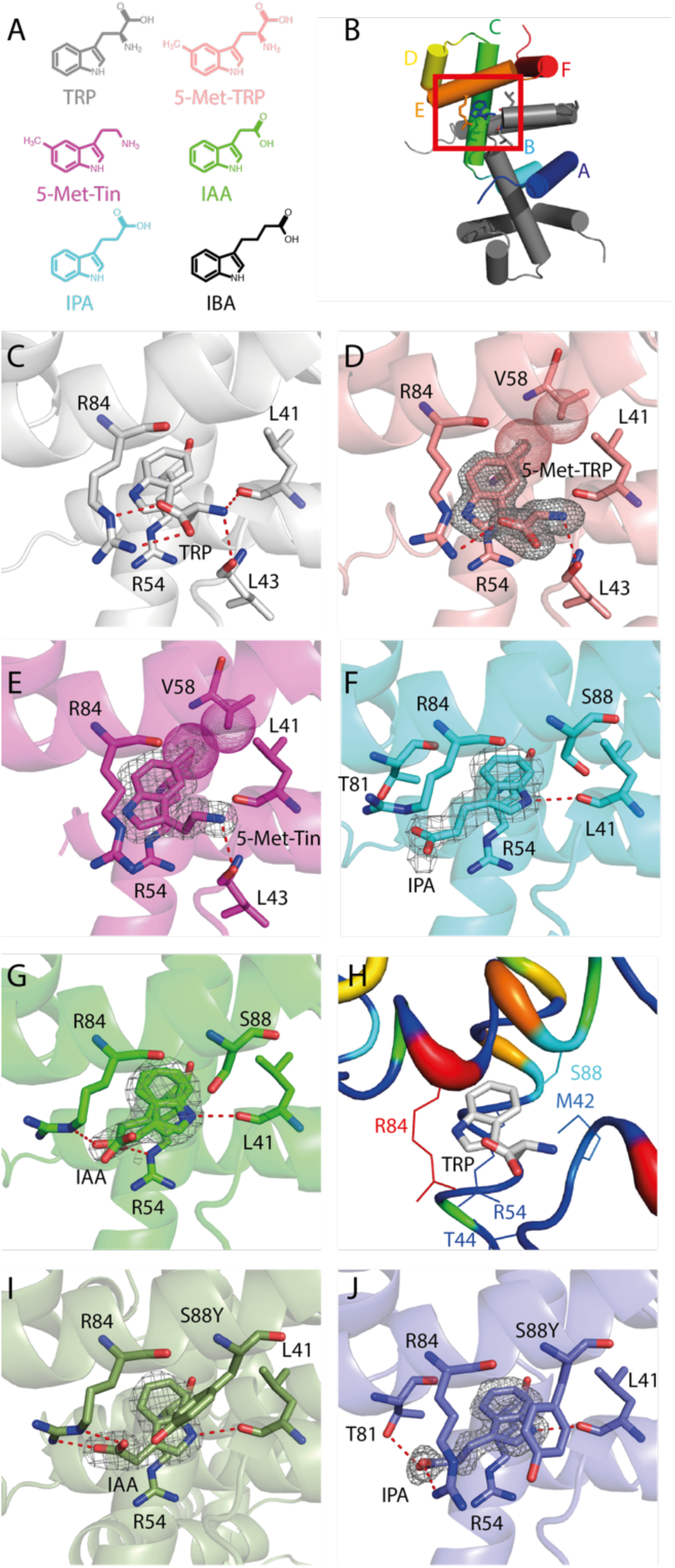
Diverse ligand binding in TrpR. (A) Ligands studied. (B) Structure of TrpR with helices labeled by letters. (C–G) TrpR bound to the native ligand TRP, the co-repressors 5-methyl-tryptophan and 5-methyl-tryptamine, and the pseudo-repressors IPA and IAA. (H) Binding pocket colored by rmsd of the former five structures (red = high, blue = low). (I–J) TrpR-S88Y with IAA and IPA. Shown are magnifications as indicated in (B). Ligand *2F*_*O*_*-F*_*C*_ density is contoured at 1σ. Important interactions are indicated by red dashed lines (electrostatic) and spheres (van-der-Waals).

## Results

The binding pocket of TrpR can be separated into two sections: (i) the hydrophobic back harboring the indole ring and (ii) the frontal part that is open to the solvent and accommodates the 3’ extensions to the indole. In particular, Arg_84_ can interact with the 3’ substituents and acts as a latch across the pocket. Accordingly, our analysis of the binding modes of TrpR to non-native ligands focused on three prominent chemical alterations to TRP: (i) hydrophobic additions to the indole ring to probe the back of the binding pocket, (ii) a decarboxylated and (iii) a deaminated Cα position of TRP to analyze interactions of different 3’ substituents.

### Hydrophobic packing of the indole ring

Helix C and B (of different monomers) form the back of the ligand-binding pocket. Despite the indole ring fitting tightly into this pocket, ligands with hydrophobic substituents of a certain size pointing to the back of the pocket do not impair binding^5^. The structure of 5-methyl-tryptophan (5-Met-TRP), which we solved at 1.4 Å resolution, provides an explanation: Its methyl group packs tightly between the side chains in the back of the hydrophobic pocket with only minor changes to the adjacent residues (Figure 1D). Despite the hydrophobic packing, which most likely introduces favorable van-der-Waals interactions, the overall affinity is lower than the one for tryptophan. This is most likely due to the lacking interactions with Leu_41_ as reflected in the relative loss of enthalpic contribution to binding. Moreover, the additional carbon at the back of the pocket does not alter the position of the indole ring compared to TRP suggesting that the hydrophobic pocket is a rather flexible fit and the positioning of the indole largely a product of the Arg_54_ π-stack as well as the stabilizing 3’-substituent at the entrance of the pocket. Nonetheless, the interaction of the 5-Met substituent especially with the Val_58_ sidechain might explain the unusually decreased entropic term due to lowered configurational entropy of residues at the back of the pocket.

### Decarboxylation of the 3’-substituent’s α-carbon

For TRP the 3’-substituent anchors the ligand in two ways: via a salt bridge between its carboxyl and the guanidino group of Arg_84_ and by interaction of its amino group with the carbonyls of Leu_41_ and Leu_43_. In the complex structure with the decarboxylated ligand 5-Methyl-Tryptamin (5-Met-Tin), which we solved at 1.7 Å resolution, the former interaction is missing. This results in the ligand being pushed further to the back (Figure 1E), suggesting that, if not counteracted by stabilization at the 3’-substituent, a “deeper” positioning of the indole ring is adopted (Supp. Figure S2). The ethylamine substituent itself is positioned similar to the corresponding atoms in TRP resulting in contacts with the backbone carbonyls of the leucine residues. However, the lacking stabilization by the carboxyl group is reflected in a two-fold loss in affinity compared to TRP due to less favorable enthalpic contributions, which can be compensated only to some extend by a higher configurational entropy of Arg_84_.

### Deamination of the 3’-substituent’s α-carbon

The opposite modification is a deamination of the 3’-substituent’s α-carbon, which yields indole-3-propionic acid (IPA). IPA is bound in a strikingly different way with the indole ring flipped by 180° and the 3’ position now pointing towards Arg_84_ (Figure 1F). The position was already suggested by NMR^12,13^ and difference-diffraction-calculations^10^ but is shown here for the first time with structural detail (2.1 Å resolution). Despite this flip the indole is congruent to its position in TRP and most of the binding pocket is unchanged. The only exception is Arg_84_, which adopts a completely different state to the one in the TRP bound structure. Its guanidino group is now pointing away from the pocket. Nonetheless, a clear electrostatic interaction with the carboxyl-group of IPA is maintained. A similar position for Arg_84_ is observed for a structure bound to Indole-3-acetic acid (IAA), which is one carbon shorter, solved to 2.0 Å resolution (Figure 1G). This indicates that the arginine position is related to the position of the carboxyl group in the flipped ligand. The IAA bound structure has no clear density for the carboxyl, pointing towards a less stable interaction. This is reflected in the poor binding affinity of TrpR towards IAA compared to IPA. The even more elongated Indole-3-butyric acid (IBA) is bound with a similarly high affinity as IPA. The increase in affinity of IPA and IBA compared to IAA might also be due to the favorable desolvation of IPA and IBA compared to IAA (inferred by solubility: IAA > IPA > IBA).

In contrast to 5-Met-TRP and 5-Met-Tin the ligands IPA and IAA are strong pseudo-repressors precluding interaction with the operator DNA sequence (> 300nM, ^6,14^). This is caused by the following: (i) The repositioned Arg_84_ guanidino group is in the wrong position for an interaction with the A+2 DNA-phosphate as can be seen by superimposing the deaminated structures with the TRP bound DNA-complex-structure^15^ (Supp. Figure S3). (ii) At least in the case of IPA the propionic acid substituent actively interferes with the DNA (A+1 phosphate) both sterically and due to electrostatic repulsion. (iii) Finally, the reading-head could be distorted as indicated by poorly resolved density indicative of structural flexibility. This is similar, but less pronounced, as in the apo-structure where Arg_84_ of the reading-head collapsed into the binding pocket ^9^.

### “Indole-based” promiscuity

The 180° flipped position of the deaminated ligands showcases the potential of TrpR in tolerating various 3’-substitutions to its ligands. The reason is that in the flipped position such substituents are pointing towards the pocket opening, which provides more spatial freedom. Moreover, Arg_84_ provides high internal flexibility and is part of the reading-head region, which adds flexibility on the backbone level and allows Arg_84_ to interact favorably with different 3’-substituted ligands (Figure 1H and Supp. Figure S4). This flexibility stands in contrast to the rigidity of stabilization of the indole ring itself. All residues involved are part of the inner core of TrpR: the hydrophobic pocket residues Leu_41_, Ile_57_, Val_58_, the π-stacking Arg_54_ and its interaction partner Glu_47_ belong to helices that form the rigid center of TrpR. Not surprisingly, none of our structures showed a binding mode that perturbs this core. This contrast between the indole-stabilizing core, tolerating only small hydrophobic substitutions, and the 3’-substituent accommodating flexible region explains TrpRs’ “indole-based” promiscuity. Such a division between a stabilizing and a variable part can also be found in other binders with broad ligand spectra. For example, the periplasmic binding protein PotF shows affinity to different polyamines^3^. In this case an invariable aromatic box stabilizes the hydrophobic middle part of the ligands.

### Design of TrpR ligand specificity

The discussed features of a binding pocket are important to consider in the rational design of ligand specificity based on promiscuous pockets. Design strategies that are in clear conflict with the “core” part of the binding pocket are unlikely to succeed since they are in conflict with the overall topology of the protein.

The 180° flipped position of the deaminated ligands is highly discriminative and, moreover, the low affinity of IAA is the most different compared to the native TRP. This provides a promising vantage point for an exemplary rational design, which prompted us to change the TrpR binding pocket towards IAA affinity. Based on a structural overlay of TrpR bound to TRP and to IAA, alterations to the sidechain of residue 88 were identified most likely to prohibit TRP binding and simultaneously stabilize the 180° flipped position of IAA. Thus, with the rationale of blocking the TRP orientation with a bulky residue and simultaneously providing hydrogen bond capabilities we mutated Ser_88_ to Tyr. The mutant TrpR-S88Y exhibits an entirely flipped specificity: IAA binding is improved ∼10 fold compared to TrpR while TRP binding is drastically reduced in the mutant.

The crystal structure of the mutant in complex with IAA (solved to 1.9 Å resolution) compared to TrpR with TRP bound confirms our rationale and shows that Tyr_88_ effectively blocks the position for the amino acid group of TRP and, thus, prohibits its interaction with the backbone carbonyl groups of Leu_40_ and Leu_43_ (Figure 1 C and I). Moreover, the fact that we could not detect any binding for TRP in this variant suggests that an alternative - most likely flipped –orientation of TRP is not possible. In contrast, for IAA binding the tyrosine is highly favorable for several reasons (Figure 1I): i) the hydroxyl group can form a hydrogen bond with the carboxyl group of IAA, ii) the phenyl ring tightly packs against the pyrrole of the indole and iii) the bulky sidechain excludes water from the pocket thereby reducing desolvation cost. Moreover, the IAA tyrosine interaction poises the carboxyl of IAA optimally for interaction with the guanidino group of Arg_84_ as reflected by a gain in enthalpic contribution of TrpR-S88Y compared to TrpR (Table 1).

Next, we clarified how the tyrosine in the binding pocket acts on the already well binding ligand IPA. Binding data shows a comparable improvement in affinity. Interestingly, the crystal structure in complex with IPA (at 1.2 Å resolution) points to slightly different factors for the improved affinity (Figure 1J). Above points ii) and iii) also apply for IPA, however, an interaction between the carboxyl group of IPA and Tyr_88_ cannot be established. Moreover, the favorable interaction between the carboxyl group of IPA and the guanidino group of Arg_84_, observed in the wild type, appears less effective and the interaction with Thr_81_, as seen in the wild type is completely abolished. Instead, Arg_84_ adopts a new conformation different from all other structures, which results in tight packing around the ligand now almost closing the pocket. These favorable interactions are reflected by a strong gain in enthalpy whose effect is only slightly reduced by a rise of the entropic penalty.

## Discussion

Our structural work shows with high detail how different ligands can be accommodated in the same binding pocket. Such comprehensive sets of structural data on different binding modes together with thermodynamic data provide an indispensable study case and starting point for *in silico* analyses since it allows studying discriminating effects based on small differences in the substituents. Our basic exemplary design shows how structural knowledge on ligand binding promiscuity can be exploited for (semi-) rational design by enhancing an existing poor affinity and diminishing it for the native ligand. In this case the increased affinity to IAA, as a ligand of extreme importance for plant biology, is a foothold to e.g. engineer molecular sensors or IAA scavengers. Moreover, such mutational experiments improve our general understanding of how molecular selectivity can be established. For TrpR we envision affinity towards indole-derivatives of broader interest can be designed, e.g. the neurotransmitter serotonin is closely related to 5-Methyl-Tryptamin, with the 5-Methyl group substituted by a hydroxyl-group or the hormone melatonin, which also shares the indole scaffold. Despite its challenges for the reasons described above, redesign of the back of the hydrophobic pocket making it more amenable for polar groups might introduce even more affinities paving the way for a range of molecular sensors.

## Experimental procedure

### Ligands

5-methyl-tryptophan (order no. M0534), 5-methyl-tryptamine (order no. 134228), indole-3-acetic acid (order no. i2886), indole-3-propionic acid (order no. 57400) and indole-3-butyric acid (order no. i5386) have been obtained from Sigma-Aldrich. All ligands have been dissolved in 50mM Tris / 300mM NaCl pH8 buffer with if necessary 1% dimethyl-sulfoxide (DMSO). For 5-methyl-tryptophan a mixture of L- and D-stereoisomers has been used. Despite expected preferential binding (the K_D_ of D-tryptophan is ∼ 25 times higher compared to L-tryptophan) this might explain the poorly resolved densities for the amino acid of 5-methly-tryptophan as well as a high error in the calorimetry.

### Protein purification

After subcloning to pET21b(+) TrpR-Wt was expressed in *E. coli* BL21 (DE3) and purified via a Ni-IMAC column and a subsequent Superdex-S75 gel filtration column. All purification steps and measurements have been based on the above 50 mM Tris / 300 mM NaCl pH 8 buffer.

### Crystallization

Crystals of TrpR with different ligands were obtained by standard vapor diffusion. The crystals were flash cooled in liquid nitrogen. Data of single crystals was collected at the synchrotron beamline PXII (Swiss Light Source, Villigen PSI, Switzerland) at 100 K and 0.5° images were recorded on a Pilatus 6 M detector. The structure with 5-Met-TRP was recorded at BL14.2 (BESSY II electron storage ring operated by the Helmholtz-Zentrum Berlin) at 100 K and 0.1° images were recorded on a Pilatus 2M detector. Data was indexed, integrated and scaled with XDS and converted with XDSCONV^16^. Molecular replacement was performed with Phenix using the coordinates of TrpR-Wt (PDB 1wrp)^17^ as search model. Model building was performed with the program Coot^18^ and refinement with Phenix^19^. Details on crystallization conditions and data and refinement statistics for all structures are summarized in (Table S1).

### Isothermal Titration Calorimetry (ITC)

ITC was performed using a nanoITC LV (TA Instruments). The protein concentration was adjusted, depending on the expected K_D_, between 500 µM and 2 mM. Accordingly, 8-10 fold molar concentrated ligand solutions were prepared using the above buffer containing 1% DMSO. Measurements were performed at 25°C with a stirring speed of 300 rpm and spacing of 250 s between injections. The data was analyzed using the NanoAnalyze program (TA Instruments). Binding data was derived from sigmoidal fits based on a one-site binding model from three measurements for each variant. Heat-of-dilution baselines for the ligands alone have been subtracted from the experimental data as described in ^20^.

### Mutagenesis

Mutagenesis was performed on the pET21b(+)-TrpR-Wt plasmid using the QuikChange technique (Stratagene) following the manufacturer’s protocol.

## Acknowledgements

We thank the beamline staff at the Swiss Light Source and at BESSY for support, and Michael Weyand for helpful discussions on crystallographic issues. We acknowledge financial support and allocation of synchrotron beamtime by HZB as well as funding by the Deutsche Forschungsgemeinschaft (grant HO 4022/2-3).

## Conflict of interests

The authors declare that they have no conflicts of interest with the contents of this article.

## Author contributions

ACS and SS collected and processed structural data, SS collected ITC data, OH-S and GJ provided TrpR-S88Y variant, ACS, SS and BH analyzed all data, ACS and BH wrote the manuscript.

## Supporting information

Experimental methods, structural visualization of TrpR interactions and data collection and refinement statistics.

