## Supplementary material for "Ligand promiscuity in the tryptophan repressor – from structural understanding towards rational design": Suppl. Figures

### Supplementary Figures:

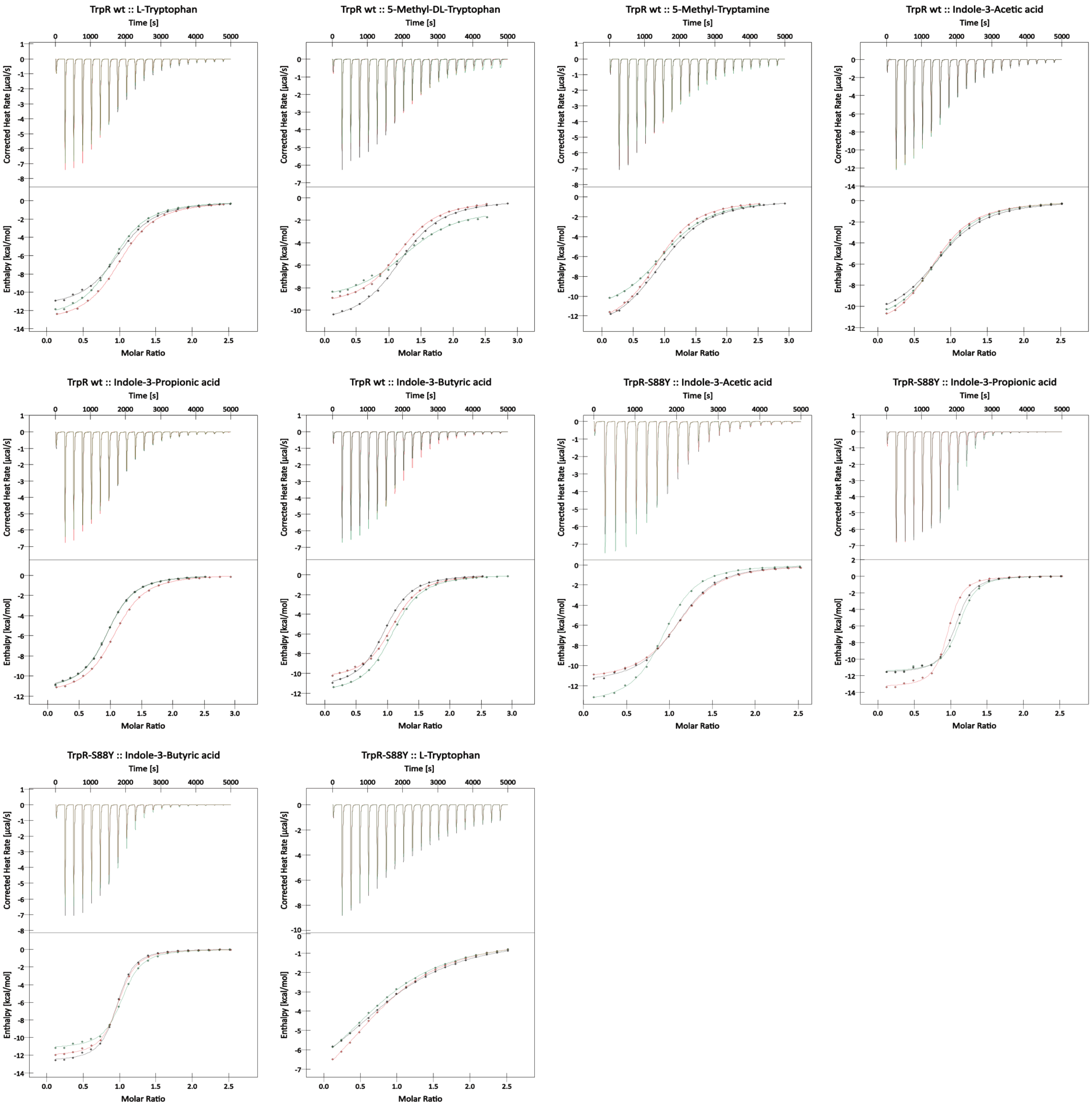


FIGURE S1**:** Analysis of binding to TrpR and the mutant TrpR-S88Y by ITC. Each plot shows three independent measurements.

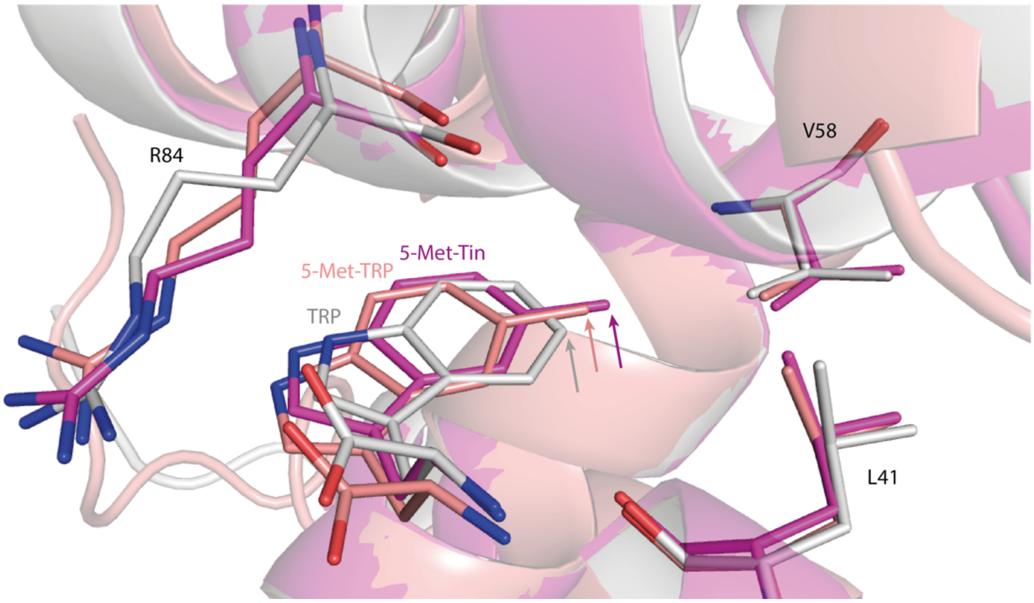


FIGURE S2**:** Packing of TRP (gray), 5-Met-TRP (salmon) and 5-Met-Tin (magenta) in the back of the binding pocket. The lacking stabilization at the 3`-substituent and the favorable packing of the methyl group is reflected in 5-Met-Tin being buried deepest in the pocket (arrows).

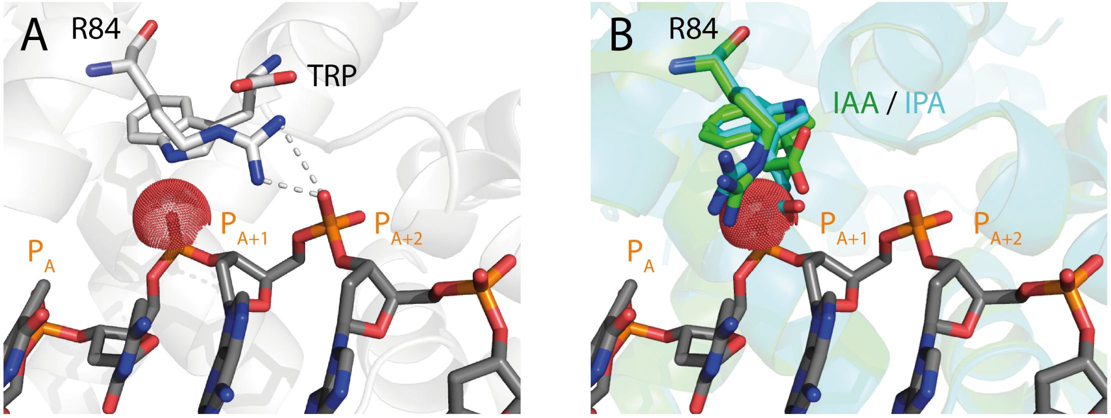


FIGURE S3**:** Interaction of TrpR with the Operator region of DNA with the native ligand tryptophan (A) and the pseudo-repressors (B) Indole-3-acetic acid (IAA) and Indole-3-propionic acid (IPA).

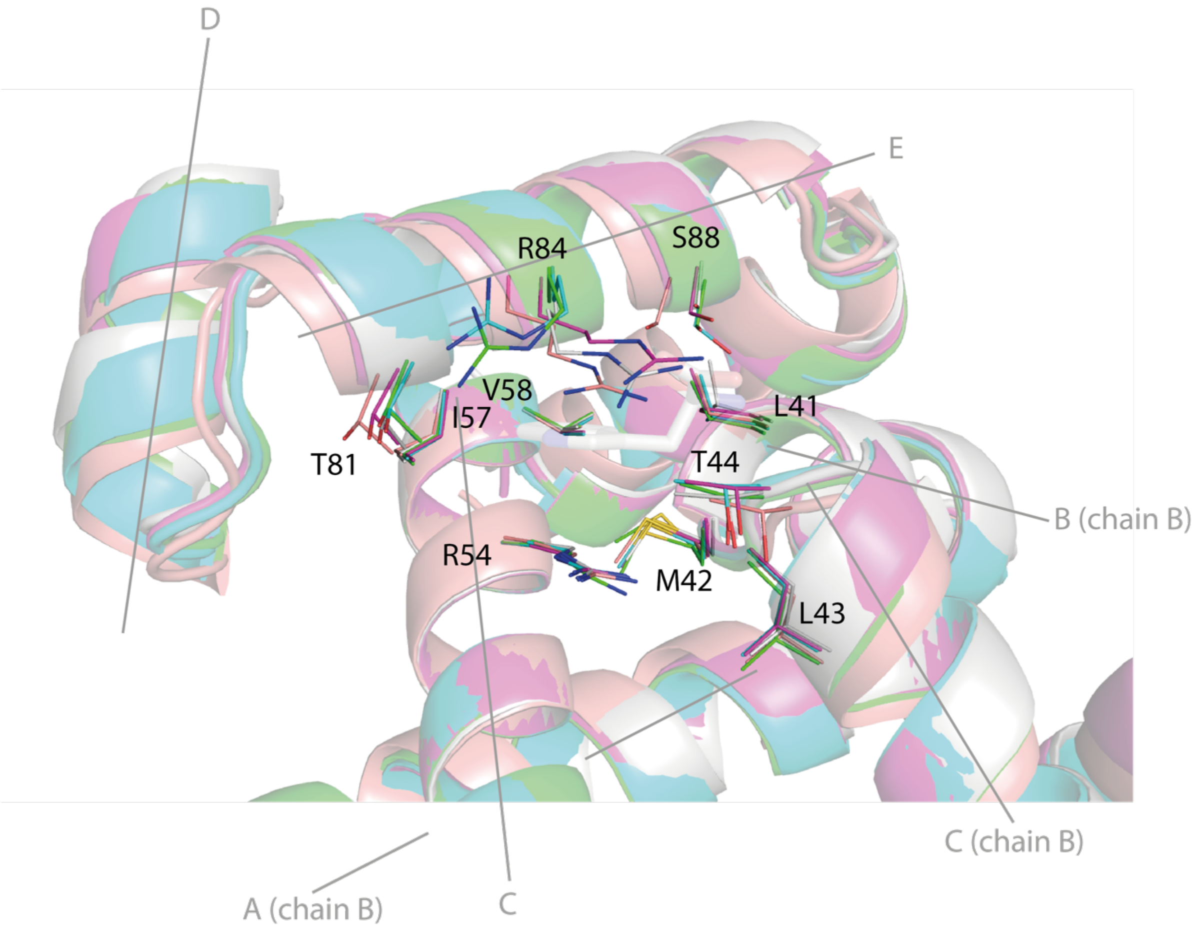


FIGURE S4**:** Superimposition (helix C) of TrpR binding different ligands (coloring as in Figure 1). Residues surrounding the ligand (for clarity only TRP shown here with transparence) shown as sticks; helical axes are indicated by letters. It is apparent that residues of the "central core" (helix B and C) like L41, M42, L43, R54, I57 and V58 show a low flexibility while residues of the "reading head region" (helix D and E) such as T81, R84 and S88 show more flexibility when accomodating the chemical nature of the different ligands. This is further refelected in the backbone variability of this region.

TABLE S1: Data collection and refinement statistics (molecular replacement) for structures elucidated in this study. Values for highest resolution shell are shown in brackets. The structures for TrpR :: IAA (6EJW) and TrpR-S88Y :: IAA (6EJZ) have been elucidated in another study.

| | TrpR :: 5-Met-TRP (6F9K) | TrpR :: 5-Met-Tryptamine (6F7G) | TrpR :: IPA (6F7F) | TrpR-S88Y :: IPA (6FAL) |
|---|---|---|---|---|
| **wavelength** | 0.92 | 1.00 | 1.00 | 0.98 |
| **Resolution range** | 20.88 - 1.41 (1.45 - 1.40) | 33.10 - 1.66 (1.72 - 1.66) | 40.92 - 2.13 (2.20 - 2.13) | 35.87 - 1.20 (1.24 - 1.20) |
| **Space group** | P 43 2 2 | P 1 21 1 | P 43 | P 1 21 1 |
| **Unit cell** | 53.15 53.15 90.83 90 90 90 | 35.35 55.55 66.99 90 98.82 90 | 81.84 81.84 70.57 90 90 90 | 36.07 45.25 60.67 90 104.02 90 |
| **Total reflections** | 222964 (21911) | 198258 (16583) | 341121 (24524) | 363609 (28813) |
| **Unique reflections** | 26306 (2544) | 30265 (2707) | 26212 (2523) | 59053 (5462) |
| **Multiplicity** | 8.50 (8.60) | 6.60 (6.10) | 13.00 (9.50) | 6.20 (5.30) |
| **Completeness (%)** | 99.53 (98.91) | 96.71 (88.58) | 99.52 (95.53) | 98.96 (92.10) |
| **Mean I/sigma(I)** | 14.94 (0.43) | 15.44 (3.76) | 12.38 (2.59) | 18.89 (6.18) |
| **Wilson B-factor** | 21.58 | 16.38 | 26.70 | 11.37 |
| **R-merge** | 0.08 (4.55) | 0.24 (0.32) | 0.17 (0.99) | 0.06 (0.25) |
| **R-meas** | 0.08 (4.84) | 0.26 (0.36) | 0.19 (1.04) | 0.07 (0.28) |
| **R-pim** | 0.03 (1.63) | 0.10 (0.14) | 0.05 (0.32) | 0.03 (0.12) |
| **CC1/2** | 1.00 (0.24) | 0.84 (0.96) | 1.00 (0.78) | 1.00 (0.95) |
| **CC*** | 1.00 (0.62) | 0.95 (0.99) | 1.00 (0.94) | 1.00 (0.99) |
| **Reflections used in refinement** | 26294 (2544) | 29657 (2707) | 26148 (2523) | 59043 (5459) |
| **Reflections used for R-free** | 1315 (127) | 1481 (136) | 1308 (126) | 2953 (273) |
| **R-work** | 0.19 (0.46) | 0.19 (0.26) | 0.17 (0.26) | 0.16 (0.21) |
| **R-free** | 0.20 (0.46) | 0.23 (0.29) | 0.23 (0.32) | 0.18 (0.22) |
| **CC(work)** | 0.96 (0.55) | 0.95 (0.93) | 0.97 (0.81) | 0.95 (0.91) |
| **CC(free)** | 0.96 (0.43) | 0.93 (0.88) | 0.93 (0.68) | 0.94 (0.90) |
| **Number of non-hydrogen atoms** | 1027 | 1995 | 3686 | 1870 |
| **macromolecules** | 877 | 1623 | 3306 | 1617 |
| **ligands** | 84 | 26 | 56 | 28 |
| **solvent** | 66 | 346 | 324 | 225 |
